# Seeking the “beauty center” in the brain: A meta-analysis of fMRI studies of beautiful human faces and visual art

**DOI:** 10.1101/081539

**Authors:** Chuan-Peng Hu, Yi Huang, Simon B. Eickhoff, Kaiping Peng, Jie Sui

## Abstract

The existence of a common beauty is a long-standing debate in philosophy and related disciplines. In the last two decades, cognitive neuroscientists have sought to elucidate this issue by exploring the common neural basis of the experience of beauty. Still, empirical evidence for such common neural basis of different forms of beauty is not conclusive. To address this question, we performed an activation likelihood estimation (ALE) meta-analysis on the existing neuroimaging studies of beauty appreciation of faces and visual art by non-expert adults (49 studies, 982 participants, meta-data are available at https://osf.io/s9xds/). We observed that perceiving these two forms of beauty activated distinct brain regions: while the beauty of faces convergently activated the left ventral striatum, the beauty of visual art convergently activated the anterior medial prefrontal cortex (aMPFC). However, a conjunction analysis failed to reveal any common brain regions for the beauty of visual art and faces. The implications of these results are discussed.

## 1. Introduction

The nature of beauty is a long-standing topic in philosophy. For example, philosophers, such as Plato, suggested that there is a “common and abstract beauty” that is independent of various concrete forms of beautiful things (Allen, 1993), and David Hume asserted a common basis for evaluating beautiful objects (Shimamura, 2012). Other scholars (e.g., Kubovy (2000)), however, have opposed this common theory by suggesting that “beauty is in the eye of the beholder,” emphasizing the role of individuals’ experiences in appreciating beautiful objects in different circumstances and refuting the assertion that there are common underpinnings to different experience of beauty.

Though these debates lie within the scope of philosophy for centuries, recently, cognitive neuroscientists and psychologists have begun to address the scientific basis of aesthetic responses to beauty via experimental approaches (e.g. (Aharon et al., 2001; Chatterjee and Vartanian, 2014; Pearce et al., 2016)). However, after two decades of investigation, it remains unclear whether there is a common neural basis for different forms of beauty (Conway and Rehding, 2013; Ward, 2015). To address this question, we synthesized the empirical studies by conducting activation likelihood estimation (ALE) meta-analyses on neuroimaging studies that explored the neural basis of beauty.

### 1.1. What is the experience of beauty?

Currently, there is no widely accepted definition for the experience of beauty. The experience of beauty was too often used together with aesthetic experiences or the appreciation of art in neuroaesthetic studies, even though they are theoretically different (Pearce et al., 2016). This conceptual ambiguity not only hampers interdisciplinary communication but also makes the cross-study comparisons difficult (Bergeron and Lopes, 2012; Conway and Rehding, 2013). Therefore, it is necessary first to offer an operational definition of beauty.

To synthesize previous neuroimaging studies on beauty, we defined the experience of beauty as a *pleasurable aesthetic experience* that is the outcome of the processing of aesthetic objects (including both art and non-art objects). This “subjectivist’s” operationalized definition of beauty has two features: First, the experience of beauty is typically accompanied by positive emotion, especially pleasure (Blood and Zatorre, 2001; Vartanian and Goel, 2004); Second, the experience of beauty as elicited by aesthetic stimuli and could be accompanied by some aesthetic emotions (Menninghaus et al., 2019). This relatively simple and broad definition of beauty can distinguish the experience of beauty from (i) other aesthetic emotions, e.g., negative aesthetic emotions (Bergeron and Lopes, 2012; Menninghaus et al., 2019; Silvia, 2009), (ii) the general pleasure, which can be elicited by non-aesthetic stimuli, e.g., food, sex (Berridge and Kringelbach, 2013), and moral behaviors (Diessner et al., 2008), and (iii) some complex emotions (e.g., “awe”, “sublime” or “being moved”), which may or may not accompany the experience of beauty. Also, this definition is consistent with the conceptual framework proposed by Pearce et al. (2016), in which the cognitive neuroscience of beauty is a sub-field of cognitive neuroscience of aesthetics.

As in previous empirical studies, we further assume that the experience of beauty can be inferred from subjective ratings of beauty, i.e., the experience of viewing a preferred object or an object with high beauty/attractiveness rating has more experience of beauty than viewing a not-preferred object or an object with low beauty rating. This operationalization provides us with specific criteria in selecting empirical studies. That is, brain imaging studies of beauty are those that directly compared brain activities elicited by beautiful stimuli to brain activities elicited by physically similar but not-beautiful stimuli, or those that correlated the brain activities with beauty ratings or preference.

Note that we are aware that this subjective definition of beauty, as pointed by (Conway and Rehding, 2013), may be fundamentally flawed if we are to seek the neural basis of a universally agreed beauty. However, this definition can suit our purpose, i.e., synthesizing the current neuroaesthetic literature to search the “beauty center” in the brain. We assumed that if there are convergent results based on all laboratory studies in which subjective definitions were used, the results would at least suggest a common neural basis for the subjective experience of beauty.

### 1.2. Is there a common neural basis for the experience of beauty?

As pointed out by Pearce et al. (2016), from the computational view of cognition (Marr, 2010), brain imaging studies of beauty are mainly addressing the implementation level of experience of beauty, less on algorithmic-representational or computational levels. Thus, one way to define the common neurobiological implementation of different forms of beauty is testing the existence of overlapping brain regions in processing different types of information (e.g., beautiful faces vs. beautiful visual art). This overlap-based approach has been widely-used to searching the common neural basis of different cognitive processes. For instance, recent studies have shown that the right inferior parietal lobule engaged in processing spatial, temporal and social distance (Parkinson et al., 2014), the left intraparietal sulcus was associated with processing both perceptually salient and socially salient stimuli (Sui et al., 2015), and the dorsal anterior cingulate cortex and the left anterior insula involved in processing both the psychological and physical selves (Hu et al., 2016). In this sense, a common neural basis for different forms of beauty can be identified if there are common regions activated by experiencing different forms of beauty.

A few studies have explored the common neural basis of beauty and suggested that there is a common neural basis for different forms of beauty. For example, Ishizu and Zeki (2011) used the conjunction analysis to find the common the brain regions that responded to both paintings and music, and they revealed that the neural basis of the experience of beauty across modalities was the medial orbitofrontal cortex (mOFC). Similarly, Pegors et al. (2015), using multi-voxel pattern analysis, compared the activation patterns of vmPFC elicited by beautiful places with the pattern of vmPFC elicited by beautiful faces. They found that the vmPFC activation pattern predicted both forms of beauty, suggesting that vmPFC not only involves in the processing of experience of different forms of beauty. Brown et al. (2011) conducted a meta-analysis using aesthetic appraisal as article inclusion criteria and looked for the overlapping voxels across the meta-analytical results from visual, audition, gustation, and olfaction studies. They found that the insula might be the common neural basis. Overall, these results were consistent with the implicit theory of common beauty that beauty is equal to reward (Aharon et al., 2001; Bray and O’Doherty, 2007; Chatterjee and Vartanian, 2014; Cloutier et al., 2008; Lacey et al., 2011; Liang et al., 2010; Smith et al., 2014).

Differing from these previous meta-analysis (Brown et al., 2011), the current meta-analyses focused on the neural basis of beauty instead of aesthetic appraisal in general. This strategy was used for two reasons. First, the results of single neuroimaging study may suffer from low statistical power (Button et al., 2013) and high false-positive rates (Carp, 2012; Eklund et al., 2016; Wager et al., 2007). Coordinate-based meta-analysis (Wager et al., 2007) is one solution to examine the cross-study convergence and the activation likelihood estimation (ALE), a widely-used coordinate-based meta-analysis technique, can accommodate the spatial uncertainty of activation data and allows researchers to form statistically defensible conclusions (Eickhoff et al., 2016; Fox et al., 2014; Laird et al., 2011). Second, the heterogeneity of fMRI studies (Liu et al., 2018; Müller et al., 2018) makes cross-study comparison difficult. That is, previous studies used aesthetic stimuli varied in a great degree, many of them did not target the neural basis of beauty. For example, some studies investigated how aesthetic judgments were made (Bzdok et al., 2012; Ishizu and Zeki, 2013), others explored how visual art was processed in the brain (Mizokami et al., 2014). It became increasingly clear that experience of beauty is not equal to the visual processing of art or aesthetic judgment itself (Pearce et al., 2016). Therefore, synthesizing studies that specifically focused beauty would provide a more direct answer to the inquiry about the common neural basis of beauty.

### 1.3. The current study

Considering the above concerns, we conducted ALE meta-analyses of fMRI studies of beauty and compared the neural activities elicited by the beauty of faces and the beauty of the visual art (includes paintings, visual patterns, architectures, and dance).

We selected the beauty of visual art and the beauty of faces for two reasons. First, they are the two most intensively studied beautiful stimuli in laboratory settings (e.g. Chatterjee and Vartanian (2014)), thus providing enough studies for meta-analyses (Müller et al., 2018). Second, they represent two typical categories of beauty: faces are the most representative natural beauty in social life, the preference of which is shaped by both evolution (e.g., Little et al. (2011)) and environment (Germine et al., 2015); while visual art is the most representative artificial beauty, which reflects the subjective aesthetic preference of human beings. A comparison between the neural responses to these two types of beauty can provide valuable start point for the exploration of common neural bases of beauty in general.

The primary goals of the current study are (1) identifying common brain regions activated by both the beauty of visual art and faces, (2) assessing the specific brain regions activated by the beauty of faces and the beauty of visual art. To fulfill these goals, we first conducted a systematic literature search of the neuroimaging studies on the beauty of visual art and faces, then selected articles based standards derived from our definition of beauty. Based on the data extracted from 49 selected neuroimaging studies, we conducted ALE meta-analyses for the beauty of visual art and the beauty of faces separately, then we used conjunction and contrast analyses of the meta-analytical results to identify the common and distinct neural basis of the beauty of visual art and faces.

## 2. Methods

### 2.1. Literature search and study selection

Articles included in the present meta-analyses were identified based on a systematic literature search using specific terms in PubMed and the Web of Science (up to Dec 12, 2018). “Face” or “facial” were paired with “attractiveness,” “beauty” or “aesthetic” for aesthetic studies of faces; and “paintings”, “visual art”, “architecture”, or “dance” were paired with “beauty” or “aesthetic” for aesthetic studies of visual art. All terms were each combined (“AND”) with “functional magnetic resonance imaging or fMRI” or “Positron emission tomography or PET” to identify relevant functional neuroimaging studies. For complete coverage, articles were also identified from recent meta-analyses and reviews (Boccia et al., 2016; Brown et al., 2011; Bzdok et al., 2011; Kirsch et al., 2016; Mende-Siedlecki et al., 2013; Vartanian and Skov, 2014). Additional studies were identified by searching through the reference lists of studies obtained via the initial search.

In our primary analysis, we selected studies based on the following criteria:

1. Only studies reporting whole-brain analyses were included, while studies based on partial coverage or employing only region-of-interest analyses were excluded. One study was included after the author provided the whole brain analyses with the contrast of current meta-analyses interested in (Lebreton et al., 2009).
2. Articles reported results as coordinates in a standard reference frame (Talairach and Tournoux or MNI). To address problems induced by different coordinates used across the studies, we converted those coordinates published in the Talairach space to the MNI space using the Lancaster transformation (Lancaster et al., 2007).
3. Only studies with non-expert young and middle-aged adults (18-50 years old) were included; studies that included art experts were excluded if they did not report results for non-experts separately (Kirk et al., 2009a) due to the influence of expertise on aesthetic appreciation (Hekkert and Wieringen, 1996).
4. Studies investigated the aesthetic appreciation of visual art or faces. According to our operationalization of beauty, we only included studies reporting the effect of beauty or the preference of faces and visual art. We have three sub-criteria: 4a), studies using visual art or art-like stimuli or faces as stimuli; 4b), studies reporting the effect of beauty or the subjective preference for visual art or faces separately and directly, therefore, studies using visual art or faces as stimuli that did not report the effect of beauty or preference were excluded; also studies did not report the effect of faces or visual art separately were excluded; and 4c), studies that included a high-level baselines (i.e., beauty art > not-beautiful art or beautiful faces > non-beautiful faces), instead of low-level baselines (e.g., photos or resting state). Note that sub-criteria 4b and 4c were not used for the additional analyses (see Supplementary Material).

Coordinates of the peak activations of brain regions, which serves as our principal measures, were extracted from each article by searching the contrast of interest from both the main text and supplementary materials. For each experiment, only one contrast’s coordinate data was extracted. Beside the coordinates data, we also reported the number of valid participants, mean age or range of age, number of male participants, modality (fMRI or PET), stimuli, tasks, and the contrast from which the data were entered our meta-analyses (see Table 1).

**Table 1.** Overview of the studies and contrasts included in the present meta-analyses.

| Articles | Model | Subjects<br>(Male) | Mean<br>age | Stimuli | Task | selected contrast |
| --- | --- | --- | --- | --- | --- | --- |
| Abitbol et al. (2015) | fMRI | 24 (13 M) | 25 | paintings | pleasantness rating | correlation with pleasantness |
| Boccia et al. (2015) | fMRI | 20 (11 M) | 25.45 | paintings | esthetic judgment | like > dislike |
| Calvo-Merino et al (2018) | fMRI | 6 (6M) | 26 | dance videos | observation | higher 'like' score > lower score |
| Cross et al (2011) * | fMRI | 22 (13M) | 24.8 | dance videos | liking/ability judgment | movement > still position |
| Cupchik et al (2009) * | fMRI | 16 (8M) | NA | paintings | pragmatic/aesthetic viewing | aesthetic > pragmatic viewing |
| Di Dio et al (2016) * | fMRI | 19 (8M) | 27.16 | paintings | observation; aesthetic or<br>movement judgment | paintings > baseline in aesthetic<br>judgment |
| Di Dio et al (2011) a * | fMRI | 32 (16M) | 19-30 | sculpture &<br>human body | observation; aesthetic<br>judgment | canonical sculpture > real human<br>body in aesthetics judgment |
| Di Dio et al (2011) b * | fMRI | 24 (12M) | 19-30 | sculpture &<br>human body | observation; aesthetic<br>judgment | canonical sculpture > real human<br>body in aesthetics judgment |
| Di Dio et al. (2007) | fMRI | 14 (8 M) | 24.5 | sculpture | observation | beautiful > not beautiful |
| Fairhall et al (2008) * | fMRI | 12 (7M) | 25 | paintings | familiarity judgment | paintings > scrambled paintings |
| Flexas et al. (2014) | fMRI | 24 (12 M) | 23.5 | paintings | beautiful or not | beautiful > not beautiful |
| Harvey et al. (2010) | fMRI | 87 (NA) | NA | paintings | preference ratings | correlation with preference |
| Huang et al (2011) * | fMRI | 14 (8M) | 20-27 | portrait paintings | observation | authentic cue > copy cue |
| Ishizu and Zeki (2011) | fMRI | 21 (9 M) | 27.5 | paintings | beauty ratings | beautiful > (indifferent + ugly) |
| Ishizu et al (2013) * | fMRI | 21 (11M) | 28.8 | paintings | aesthetic/brightness judgment | aesthetic > brightness judgment |
| Jacobs et al. (2012) | fMRI | 18 (10 M) | 20-39 | visual textures | beauty judgment | beautiful > ugly |
| Jacobsen et al (2006) | fMRI | 15 (6M) | 25.4 | visual pattern | aesthetic/symmetry judgment | beautiful > not-beautiful judgment |
| Kawabata and Zeki (2004) | fMRI | 10 (5 M) | 20~31 | paintings | beauty ratings | beautiful > neutral |
| Kesner et al. (2018) * | fMRI | 24 (13M) | 30 | portraits | gaze direction | painterly style > linear portraits |
| Kirk et al. (2008) * | fMRI | 15 (9M) | 24.4 | photos | aesthetic ratings | correlated with aesthetics ratings |
| Kirk et al. (2009b) | fMRI | 14 (9 M) | 26.3 | paintings | aesthetic rating | correlation with aesthetics ratings |
| Lacey et al. (2011) | fMRI | 8 (4 M) | 23.1 | paintings | animacy rating | correlated with beauty |
| Lebreton et al. (2009) | fMRI | 20 (10 M) | 22.0 | paintings | pleasantness ratings | correlated with pleasantness |
| Lutz et al. (2013) * | fMRI | 20 (10M) | 35 | artworks | attractive judgment | artworks > photos |
| Mirura et al (2010) * | fMRI | 49 (38M) | 19-29 | dance videos | observation | Human > smooth robotic dance |
| Mizokami et al (2014) * | fMRI | 39 (NA) | 27.5 | paintings | aesthetic judgment | paintings > photos |
| Silveira et al. (2015a) | fMRI | 17 (8M) | 37.0 | paintings | aesthetic judgment | positive > negative aesthetic judgment |
| Silveira et al (2012) * | fMRI | 15 (7M) | 26 | paintings | affected or not | naturalistic > surrealistic |
| Silveira et al. (2015b) * | fMRI | 15 (7M) | 26.5 | paintings | emotional involvement | naturalistic > surrealistic |
| Thakral et al. (2012) | fMRI | 16 (NA) | NA | paintings | pleasant judgment | correlated with aesthetic ratings |
| Vartanian and Goel (2004) | fMRI | 12 (4 M) | 28 | paintings | preference rating | correlated with preference |
| Vartanian et al (2013) | fMRI | 18 (6M) | 23.4 | architectural space | beauty/approach-avoidance judgment | parametrically correlated with beauty ratings |
| Vessel et al. (2012) | fMRI | 16 (11 M) | 27.6 | visual arts | recommendation | most recommended > least recommended |
| Wiesmann et al (2010) * | fMRI | 12 (7M) | 24 | paintings | object recognition | paintings > scrambled |
| Zhang et al. (2016) | fMRI | 16 (9M) | 22.2 | pictograph | aesthetic judgment | beautiful > ugly pictograph |
| Zhang et al (2017) | fMRI | 19 (7M) | 21.7 | visual symbols | aesthetic judgment | beautiful > ugly pictograph |
| Aharon et al. (2001) | fMRI | 10 (10 M) | 25.2 | faces | observation | beauty > average |
| Bray and O'Doherty (2007) | fMRI | 25 (12 M) | 20.8 | faces | location discrimination | attractive > unattractive faces |
| Cartmell et al. (2014) | fMRI | 16 (7 M) | 20 | faces | Partner Selection | attractive > unattractive faces |
| Chatterjee et al. (2009) | fMRI | 13 (6 M) | 22.6 | faces | beauty/ identity ratings | correlation with beauty ratings |
| Chien et al. (2016) | fMRI | 32 (16 M) | 25.1 | faces | faces as reward cue | main effect of attractiveness |
| Cloutier et al. (2008) | fMRI | 48 (24 M) | 21.7 | faces | attractiveness judgment | increase with attractiveness |
| Cooper et al. (2012) | fMRI | 39 (20 M) | 21.44 | faces | attractiveness rating | positively related to attractiveness |
| Funayama et al. (2012) | fMRI | 42 (21M) | 18-24 | faces | choose one face from two | chosen for partners > control |
| Iaria et al. (2008) | fMRI | 11 (5 M) | 24.09 | faces | attractiveness rating | attractive > unattractive faces |
| Ito et al. (2014) | fMRI | 12 (4M) | 19.2 | generated faces | social preference rating | chosen > unchosen |
| Ito et al. (2015) | fMRI | 28 (14 M) | 21.6 | faces | passive viewing | preferred > non-preferred |
| Ito et al., (2016) * | 32 | 32 (16M) | 21.3 | faces | pleasantness rating | correlated with pleasantness |
| Kedia et al (2014) * | 25 | 25 (0M) | 23.46 | female faces | beauty judgement | beauty > distance judgment |
| Kim et al. (2007) | fMRI | 25 (13 M) | 20-45 | faces | ratings | correlated with attractiveness<br>(exclude preference) |
| Kocsor et al. (2013) | fMRI | 16 (8 M) | 25 | faces | face discrimination | attractive > unattractive faces |
| Liang et al. (2010) | fMRI | 17 (8 M) | 26.5 | faces | passive viewing | correlated with attractiveness |
| Martín-Loeches et al. (2014) | fMRI | 20 (10M) | 21.3 | faces | beauty judgment | beautiful > neutral faces |
| McGlone et al. (2013) | fMRI | 16 (0 M) | 23 | faces | attractiveness rating | attractive > unattractive faces |
| Nakamura et al (1998) * | PET | 6 (6M) | 19-25 | face | attractiveness rating | attractiveness > emotion judgment |
| O'Doherty et al. (2003) | fMRI | 25 (13 M) | 23.8 | faces | gender judgment | high > low attractiveness |
| Pegors et al. (2015) | fMRI | 28 (14 M) | 22.5 | faces | attractiveness rating | correlated with face attractiveness |
| Shen et al. (2016) | fMRI | 36 (19M) | 23.57 | faces | attractiveness rating | correlated with attractiveness |
| Silveira et al (2014) a* | fMRI | 16 (16M) | 24.13 | female faces | ratings | attractive opposite-sex faces > dot |
| Silveira et al (2014) b* | fMRI | 16 (0M) | 24.64 | male faces | ratings | attractive opposite-sex faces > dot |
| Smith et al. (2010) | fMRI | 23 (23 M) | 21.8 | faces | passive viewing | attractive > unattractive faces |
| Smith et al. (2014) | fMRI | 16 (16 M) | 23 | faces | attractiveness rating | linear increase with attractiveness |
| Tsukiura and Cabeza (2011) | fMRI | 20 (0 M) | 23.4 | faces | attractiveness rating | linear increase with attractiveness |
| Turk et al (2004) | fMRI | 18 (9M) | 22 | faces | partner choice | date partner > face choosing task |
| Ueda et al. (2016) | fMRI | 36 (36M) | 25 | female faces | partner preference | attractive > unattractive |
| Ueno et al. (2014) | fMRI | 28 (14M) | 20.7 | female faces | attractiveness rating | correlated with attractiveness |
| Vartanian et al. (2013) | fMRI | 29 (14 M) | 25.1 | faces | attractiveness rating | correlated with attractiveness |
| Wang et al. (2015) | fMRI | 22 (10 M) | 21 | faces | gender judgment | beautiful face > common face |
| Winston et al. (2007) | fMRI | 15 (15 M) | 25.5 | face | attractiveness judgment | effect of attractiveness |
| Yu et al. (2013) | fMRI | 18 (9 M) | 21 | faces | attractiveness judgment | attractive faces > unattractive faces |
| Zaki et al (2011) | fMRI | 14 (14M) | 21.8 | faces | attractiveness rating | peer rating high > peer rating low |
| Zhai et al. (2010) | fMRI | 18 (10 M) | 20.8 | faces | attractiveness judgment | attractive > unattractive faces |

\* Included based on the liberal standard but not on the conservative standard.

### 2.2. Activation likelihood estimation

The revised ALE algorithm, which was implemented in Matlab code, for the coordinate-based meta-analysis of neuroimaging results (Eickhoff et al., 2009; Laird et al., 2009a; Laird et al., 2009b; Turkeltaub et al., 2002). This algorithm aims to identify areas that exhibit a convergence of reported coordinates across experiments that is higher than expected under a random spatial association. The key idea behind ALE is to treat the reported foci not as single points but rather as centers for 3D Gaussian probability distributions that capture the spatial uncertainty associated with each focus. The Full-Width Half-Maximum (FWHM) of these Gaussian functions was determined based on empirical data on the between-subject variance by the number of examined subjects per study, accommodating the notion that larger sample sizes should provide more reliable approximations of the “true” activation effect and should, therefore, be modeled by “smaller” Gaussian distributions (Eickhoff et al., 2009). Specifically, the number of subjects in the studies in our meta-analysis ranged from 6 ~ 87, with a median of 18, and the range of Full-Width Half-Maximum (FWHM) was from 8.5 mm ~ 10.94 mm (median: 9.5 mm).

The probabilities of all foci reported in a given experiment were then combined for each voxel, resulting in a modeled activation (MA) map (Turkeltaub et al., 2012). Taking the union across these MA maps yielded voxel-wise ALE scores that described the convergence of the results across experiments at each location of the brain. To distinguish ‘true’ convergence among studies from random convergence (i.e., noise), we compared ALE scores to an empirical null distribution reflecting a random spatial association among experiments. Here, a random-effects inference was invoked, focusing on the inference on the above-chance convergence among studies rather than the clustering of foci within a particular study. Computationally, deriving this null-hypothesis involved sampling a voxel at random from each of the MA maps and taking the union of these values in the same manner as performed for the (spatially contingent) voxels in the true analysis, a process that can be solved analytically (Eickhoff et al., 2012). The p-value of the “true” ALE was then given by the proportion of equal or higher values obtained under the null-distribution. The resulting non-parametric *p*-values were then thresholded at the *p* < 0.05 (cluster-level corrected for multiple-comparison; cluster-forming threshold *p* < 0.001 at voxel level) (Eickhoff et al., 2012). All significant clusters were reported, and the volume, weighted center and locations, and Z-scores at the peaks within the regions are given.

### 2.3. Contrast and conjunction analysis of individual meta-analyses

To explore the distinct and common neural basis for two forms of beauty, we further conducted contrast and conjunction analyses based on the ALE results. Differences between conditions were tested by first performing separate ALE analyses for each condition and computing the voxel-wise difference between the ensuing ALE maps. All experiments contributing to either analysis were then pooled and randomly divided into two groups of the same size as the two original sets of experiments reflecting the contrasted ALE analyses (Eickhoff et al., 2011; Rottschy et al., 2012). The ALE scores for these two randomly assembled groups were calculated, and the differences between the ALE scores were recorded for each voxel in the brain. Repeating this process 25,000 times then yielded a null-distribution of differences in ALE scores between the two conditions. The “true” difference in the ALE scores was then tested against this voxel-wise null-distribution of label-exchangeability and thresholded at a probability of *p* > 95% for true differences. The conjunction analyses used the voxel-wise minimum of each single ALE results, i.e., finding the minimum *z*-value across the two thresholded ALE results voxel-wisely (Nichols et al., 2005).

### 2.4 Data Visualization

Given that there is no golden-standard atlas for neuroimaging studies, we used probabilistic cytoarchitectonic maps (as implemented in SPM Anatomy Toolbox, the third version) (Amunts et al., 2013; Eickhoff et al., 2006; Eickhoff et al., 2007; Eickhoff et al., 2005) to assign our resulting coordinates to anatomical structures. For visualization purposes, BrainNet Viewer (Xia et al., 2013) was used.

## 3. Results

### 3.1. Studies included in the meta-analyses

49 articles were identified (20 articles for the beauty of visual arts, including 20 independent samples, 107 foci, and 295 subjects; 29 articles using attractive faces, including 29 independent sample, 183 foci, and 687 subjects). All the code for search literature, endnote files for selecting articles, and metadata for the current study are available at https://osf.io/s9xds/. See Figure 1 for the process of article selection in detail, and Table 1 for the information of selected articles.

**Figure 1.**
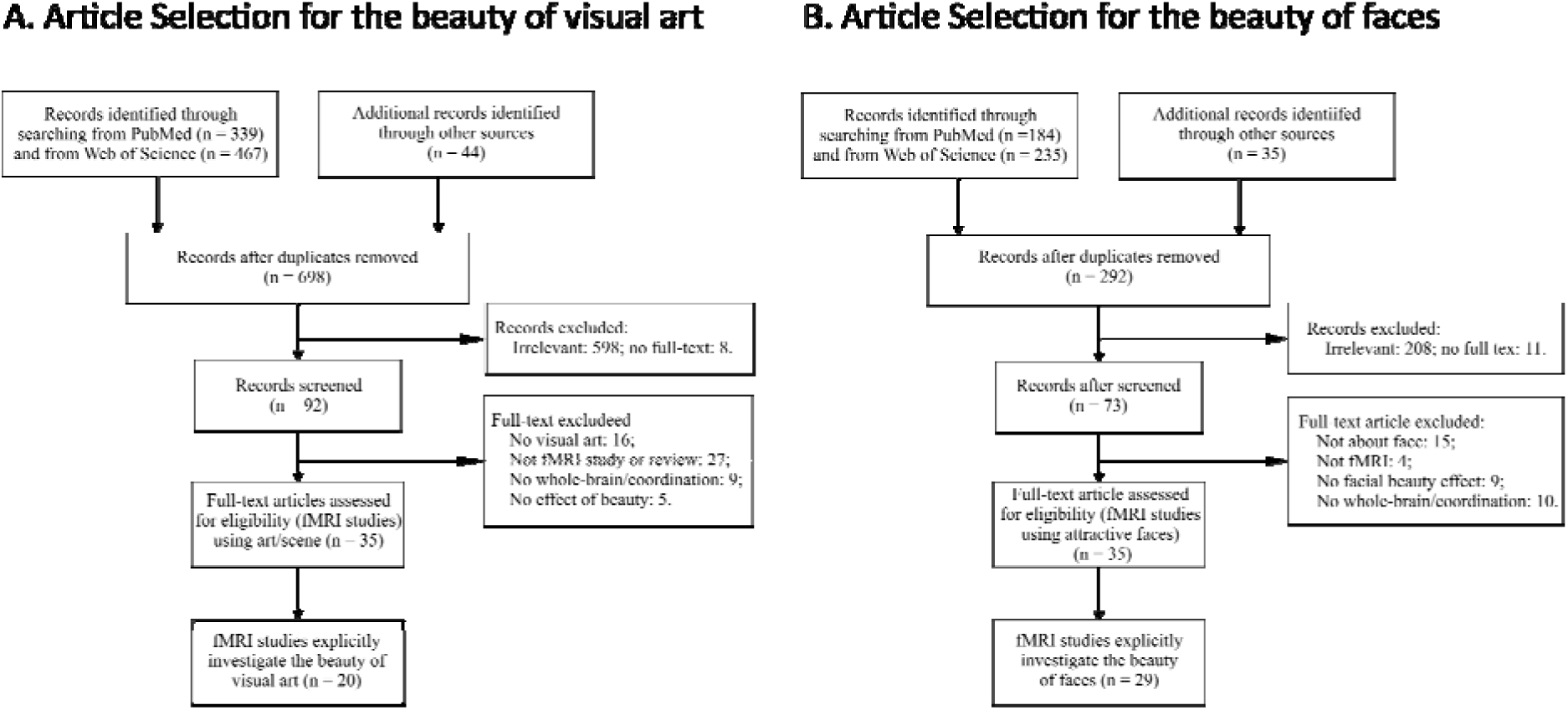
Flow diagram for literature search for the beauty of visual art (A) and the beauty of faces (B), adapted from Liberati et al. (2009).

### 3.2. ALE Meta-analyses of the aesthetic beauty of visual art and faces

Our primary analyses revealed that the frontal pole was convergently activated by the beauty of visual art. This brain region was also labeled as the anterior medial prefrontal cortex (aMPFC) in the literature (Table 2 and Figure 2A). The ALE results of the beauty of faces showed that two brain regions were more convergently activated by beautiful faces than by non-beautiful faces: the first region located in the ventromedial prefrontal cortex (vMPFC) extending to the pgACC; while the second region includes subcortical structures such as the ventral striatum and subcallosal cortex (Table 2 and Figure 2B).

**Figure 2.**
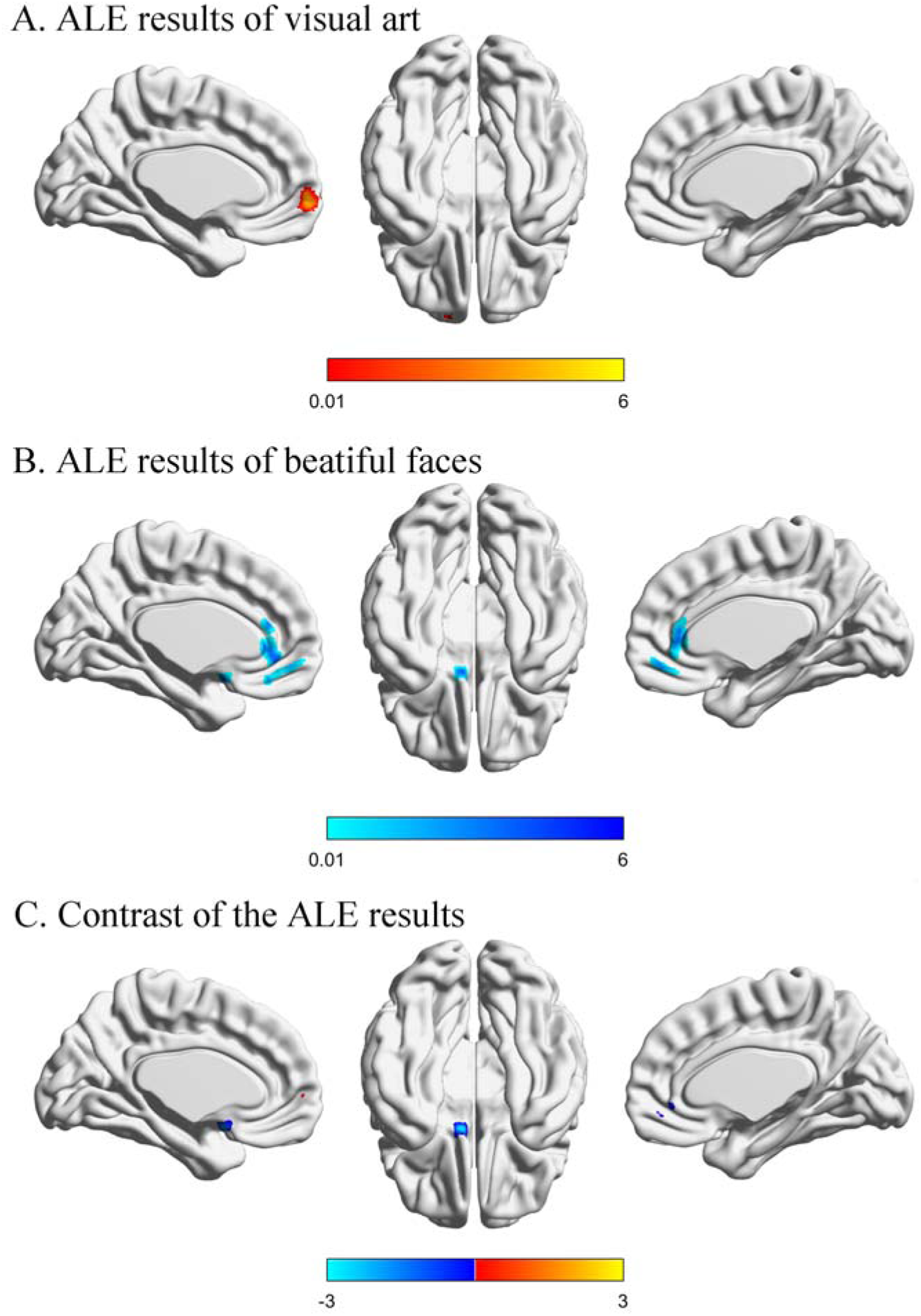
Results of the ALE meta-analysis and the contrast analysis. (A) Brain regions convergently activated more for beautiful visual art than for non-beautiful visual art; (B) brain regions convergently activated more for beautiful faces than for non-beautiful faces; (C) the results of the contrast analysis between the ALE results of beautiful visual art and beautiful faces; positive values (red) indicate greater activation for beautiful visual art than for beautiful faces, and negative values (blue) indicate greater activation for beautiful faces than for beautiful visual art.

**Table 2.** The results of the meta-analyses for beautiful visual art and beautiful faces.

| Cluster | Volume<br>(voxels) | Weighted center |  |  | Maximum<br>Z-value | Center for maximum Z-value |  |  | Macroanatomical location |
| --- | --- | --- | --- | --- | --- | --- | --- | --- | --- |
|  |  | x | y | z |  | x | y | z |  |
| *beautiful > non-beautiful visual art* |  |  |  |  |  |  |  |  |  |
| 1 | 149 | -12 | 62 | -2 | 4.33 | -12 | 62 | -2 | Frontal pole |
|  |  |  |  |  | 3.97 | -4 | 60 | -2 | Frontal pole/paracingulate gyrus |
|  |  |  |  |  | 3.97 | -6 | 60 | -2 | Frontal pole/paracingulate gyrus |
| *beautiful > non-beautiful faces* |  |  |  |  |  |  |  |  |  |
| 1 | 393 | 0 | 48 | -6 | 5.56 | 0 | 48 | -6 | Paracingulate gyrus/Frontal medial cortex |
|  |  |  |  |  | 4 | -2 | 36 | 2 | Anterior cingulate gyrus |
|  |  |  |  |  | 3.7 | 2 | 36 | 10 | Anterior cingulate gyrus |
|  |  |  |  |  | 3.2 | -4 | 36 | -16 | Frontal medial cortex/paracingulate gyrus |
| 2 | 121 | -10 | 16 | -6 | 4.69 | -10 | 16 | -6 | Left accumbens/Left caudate |
|  |  |  |  |  | 3.84 | -8 | 10 | -16 | Subcallosal cortex |
|  |  |  |  |  | 3.84 | -8 | 10 | -14 | Subcallosal cortex/left accumbens |

All peaks were assigned to the most probable brain area using the SPM Anatomy Toolbox v3

The conjunction analysis found no survival cluster. The contrast analysis further revealed that a locus within the pgACC and a locus within the left ventral striatum were more frequently activated by beautiful faces than by beautiful art, while there the left frontal pole was more activated by beautiful visual art than by beautiful faces (Table 3 and Figure 2C).

**Table 3.** Contrast and conjunction analyses of the meta-analysis results for the beauty of visual art and faces.

| Cluster | Volume | Weighted center |  |  | Maximum | Center for maximum Z-value |  |  | Macroanatomical location |
| --- | --- | --- | --- | --- | --- | --- | --- | --- | --- |
|  | (voxels) | x | y | z | Z-value | x | y | z |  |
| *the beauty of visual art > beauty of faces* |  |  |  |  |  |  |  |  |  |
| 1 | 30 | -12 | 58 | 2 | 1.98 | -12 | 58 | 2 | Frontal pole/paracingulate gyrus |
| *the beauty of faces > beauty of visual art* |  |  |  |  |  |  |  |  |  |
| 1 | 48 | 4 | 46 | -2 | 2.23 | 4 | 46 | -2 | Paracingulate gyrus/ anterior cingulate gyrus |
| 2 | 45 | -6 | 6 | -16 | 2.78 | 6 | 6 | -16 | L Accumbens/subcallosal cortex |
| *beautiful stimuli ∩ non-beautiful stimuli* |  |  |  |  |  |  |  |  |  |
| *No cluster survived* |  |  |  |  |  |  |  |  |  |

All peaks were assigned to the most probable brain area using the SPM Anatomy Toolbox v3.

## 4. Discussion

The current study sets out to find the convergent neural basis of experiencing the beauty of visual art and faces. Using standards derived from our operationalized definition of beauty, we didn’t find any overlapped brain regions between meta-analytical results of these two forms of beauty. However, different brain regions were found for different forms of beauty: the left ventral striatum and vmPFC/pgACC was more convergently activated by beautiful faces, while the aMPFC/frontal pole was more activated by beautiful visual art.

### 4.1. Is there a common neural basis for visual beauty?

Unexpectedly, our conjunction analysis failed to find any surviving clusters. This result contradicts to the neuroaesthetic literature, which suggested a common neural basis of beauty in the vmPFC. Previous fMRI studies (which were included in our meta-analysis) had found the vmPFC/mOFC was found to be involved in processing different beauties (Chatterjee and Vartanian, 2014; Ishizu and Zeki, 2011; Pegors et al., 2015). Additionally, the view that vmPFC serves as the common neural basis of beauty is in line with the literature in cognitive neuroscience: the vmPFC is a hub for integrating sensory information and abstract concepts to generate affective meaning (Delgado et al., 2016; Etkin et al., 2006; Roy et al., 2012; Sui and Humphreys, 2015), and it has been consistently activated by the positive signals (Bartra et al., 2013; Haaker et al., 2013; Kalisch et al., 2006), self-referential process (Hu et al., 2016; Northoff et al., 2006), and more importantly, value (Levy and Glimcher, 2012). By synthesizing the available studies, the present meta-analysis did not find any positive convergent results. Given the greater statistical power of meta-analysis and its capacity to address the cross-study variance, the present study may provide compelling evidence that the common neural basis of beauty does not exist.

One possible explanation of our negative results is that our meta-analysis based on a restricted criterion and didn’t include enough studies, especially for studies using visual arts. However, this explanation was unlikely because our additional meta-analyses based on a more inclusive criterion showed no surviving cluster for visual art, let alone the overlapping region between visual art and faces (see supplementary analyses and results).

The contrast between the strong theoretical prediction of and lack of empirical evidence for the common neural basis for beauty calls for more rigorous studies in the future. Recent years, the field of cognitive neuroscience has been challenged on reproducibility (Hu et al., 2018; Müller et al., 2017; Poldrack et al., 2017). As we mentioned above, small sample size (which results in low statistical power), flexibility in data analysis, and errors in implementing software, accompanied by publication bias, all threatened the reliability and reproducibility of the field (Hu et al., 2018; Poldrack et al., 2017). Direct replication, albeit very few, found that a rather pessimistic view (Boekel et al., 2015). To better test the common neural basis of beauty, researchers need to adopt new practices that have been recommended in recent year, including the standard reporting, preregistration, and greater sample size (Munafò et al., 2017). Also, as science is accumulative, further studies can be integrated into the current meta-analysis to update the results. To facilitate this process, we opened the meta-data of this meta-analysis (see Method section), future works can easily be integrated to update the knowledge about the common neural basis of beauty.

### 4.2. Unique neural basis underlying the beauty of faces and visual art

The ALE results of beautiful faces showed that beautiful faces induced greater activation in the vMPFC/pgACC and the left ventral striatum than non-beautiful faces. While the ALE results of the beauty of visual art showed convergent activations in the left frontal pole/aMPFC.

These results suggest that the vMPFC-subcortical rewarding system is engaged in processing the beauty of faces. On the one hand, previous studies have shown that these two brain regions play a critical role in processing reward (Haber and Knutson, 2010; Liu et al., 2011; Vincent et al., 1993). These findings are consistent with the view that evaluating the rewarding value is crucial for appreciating the facial beauty (Chatterjee and Vartanian, 2014; Hahn and Perrett, 2014). On the other hand, the role of the vMPFC/pgACC and the ventral striatum may differ: while the vMPFC/pgACC is coactivated with a wide range of brain structures and involved in multiple higher-level functions (de la Vega et al., 2016; Roy et al., 2012), the function of the ventral striatum mainly engaged in reward processing (Haber and Knutson, 2010; Liu et al., 2011), especially primary reward (Sescousse et al., 2013). Therefore, it is possible the facial beauty is appreciated through a ventral pathway: the ventral striatum may primarily respond to the rewarding value of faces, the reward signal, along with other information, was then integrated into the vMPFC to generate positive affections.

The frontal pole/aMPFC, which is convergent activation in appreciating the beauty of visual art, is more engaged in high-level, top-down processing (Bzdok et al., 2013). Previous studies suggested that it involves in episodic memory, decision-making, and social cognition (de la Vega et al., 2016), for example, positive evaluation (Bartra et al., 2013) and secondary reward (Sescousse et al., 2013). Therefore, we speculate that the aMPFC, when appreciating visual art, links more abstract beauty to reward, like that of secondary reward.

### 4.3. Role of the sensory cortex and hemispheric differences in processing beauty

We did not find greater activations in the sensory regions in neither meta-analyses of the beauty of faces and the beauty of visual art. For the facial beauty, there were no more activations in the fusiform face areas or other sensory cortical areas for beautiful faces than for non-beautiful faces; for the visual arts, we did not observe activation of the sensory-motor network. These results seem to contradict with previous theories about facial beauty (Chatterjee et al., 2009; Iaria et al., 2008) and art appreciating (Boccia et al., 2016; Chatterjee and Vartanian, 2014; Leder and Nadal, 2014).

One possible reason is that current meta-analyses only included the contrast between beautiful vs. non-beautiful stimuli, therefore eliminating the effect of physical feature and revealed the purer neural activities for beauty. If this is the case, the current study provided evidence that after eliminating the difference in physical features, the beauty of faces and visual arts showed different neural representations in the high-level processing.

However, that does not mean that physical features are not important in the process of beauty appreciating. Actually, the sensory network is necessary for processing the beauty of faces and visual art (Chatterjee and Vartanian, 2014). Also, the shared standard for beautiful faces (Hönekopp, 2006; Leder et al., 2016) suggests that physical features are crucial for beauty. Hence future studies are needed to examine the contribution of sensory processing to beauty appreciation.

Regarding the hemisphere differences, it seems that peak locations of both face and art beauty appear in the left hemisphere, contradict with a simplified view that “the right brain is related to imagination, (art) creativity and emotions” (Bromberger et al., 2011). However, our meta-analytic results failed to reveal hemispheric asymmetry: most clusters were near the midline of the brain. The present results, together with the previous meta-analysis on appreciating of visual art (Boccia et al., 2016), suggest that the visual beauty is processed by both hemispheres.

### 4.4. Methodological considerations

Several potential limitations of the present study should be discussed. First, the current meta-analysis methods were based on the reported peak activations, which may have two limitations. One limitation is that a large part of the spatial information was discarded. However, this limitation can be alleviated by the fact that the results derived from imaged-based meta-analysis are in good agreement with coordinate-based meta-analysis approaches (Salimi-Khorshidi et al., 2009). Another limitation is that studies didn’t report any activations for the contrast of interest will not be included. In our case, studies used the beauty appreciation task but didn’t report results of “beautiful > non-beautiful” contrast and studies did not report significant activations for this contrast (Kampe et al., 2001) were not included. However, the main concern of this limitation, together with potential publication bias in neuroimaging studies (Jennings and Horn, 2012), is the inflated false positive, which is not a concern for current meta-analysis because we didn’t found positive results for the conjunction analysis.

Second, coordinate-based meta-analysis approaches of neuroimaging studies, ALE is a representative one, use the “averaged” likelihood in common volumetric space (Wager et al., 2007); thus it might lead to false positives of convergent activation in adjacent regions across studies. This, again, is not a concern for current meta-analysis given our negative results.

Third, the meta-analyses inherently including many seems trivial but important choices, these choices bring extra flexibility in research practices. For example, the standard for inclusion may be not clear for a few papers, or results from more than one contrast are qualified with the inclusion standard. In these cases, the final choice might be arbitrary. Studies have shown that the flexibility in research practice can inflate the false positive rate (Carp, 2012; Simmons et al., 2011), this might also be true for meta-analysis (Müller et al., 2018; Müller et al., 2017). To alleviate this limitation, we adopted open and transparent practice (Schönbrodt et al., 2015) and uploaded the article selection process (as recorded in Endnote^®^ X8, Clarivate Analytics, Philadelphia, USA) and meta-data (see https://osf.io/s9xds/).

### 4.5. Conclusion

Our meta-analytic results revealed distinct neural specificities for beautiful visual art and faces, but no evidence for the common neural basis of beauty. This negative result, together with recent other negative finding from meta-analyses of neuroimaging studies (Müller et al., 2017; Nickl-Jockschat et al., 2015), suggest that, as the neuroimaging community as whole, more rigorous studies are needed to test the question of interest (Button et al., 2013; Hu et al., 2018; Poldrack et al., 2017).

## Acknowledgments

This work was supported by the Economic and Social Research Council (UK, ES/K013424/1) and by the National Natural Science Foundation of China (Project 31371017, and 31471001). The authors would like to thank Dr. Steven Brown for helpful comments on the previous versions of this manuscript.

## Author Contributions

C-P. H., K.P. designed the study. C-P.H. and S.E. performed the statistical analyses. C-P. H., Y.H., J.S analyzed the findings and wrote the manuscript. All authors reviewed the manuscript.

## Additional Information

Competing financial interests: The authors declare no competing financial interests.

