## Supplemental Materials for "Seeking the “beauty center” in the brain: A meta-analysis of fMRI studies of beautiful human faces and visual art"

### Additional analysis: Method

Two additional analyses were carried out to explore the stability of the results. First, to test the possible overlapping clusters between the meta-analytic results from the beauty of visual art and faces, we examined the unthresholded Z-maps from these two ALE analyses. More specifically, the unthresholded Z-maps of visual art and faces were masked by the thresholded images of faces and visual art, respectively, and then the peak z-value in the resulting clusters were extracted. The peak z-value can be viewed as an index of how far the current results are from significant conjunction. We carried out these analyses using the image calculator in DPABI (Yan et al., 2016).

Second, to test the stability of the primary meta-analyses, we conducted meta-analyses for the beauty of visual art and faces using a more liberal article-selection standard. More specifically, we applied a more liberal standard which included all brain imaging studies that used beautiful visual art or beautiful faces (i.e., we only applied criterion 1, 2, 3, and 4a mentioned in section 2.1). In this way, we included studies that specifically investigated the effect of beauty or used low-level baselines.

### Additional analysis: Results

The peak z-value of visual art and faces, within the clusters of thresholded maps of faces and visual art, respectively, were found. The peak z-value of visual art within vMPFC/pgACC was 3.89 (coordinates, [-4 58 -2]), within the left accumbens/subcallosal cortex was 2.87 (coordinates, [-6 18 -4]). The peak z-value of faces with the frontal pole was 3.4 (coordinates, [-2 56 -4]). These results suggest that there might be a potential overlapping in vMPFC/pgACC.

As for additional meta-analyses with a more liberal article-selection standard (see method section for more details), 70 articles were identified (35 articles using visual arts or art-like stimuli, including 36 independent samples, 210 foci, and 744 subjects; 35 articles using attractive faces, including 36 independent sample, 229 foci, and 814 subjects). The ALE analysis of the beauty of visual art did find any survival clusters. The ALE analysis of beautiful faces resulted in similar two brain regions as in primary analysis including ventromedial prefrontal cortex (vMPFC, extending to the pgACC) and ventral striatum (see Table S1). These results suggest the meta-analytical results from beautiful faces are stable, while the results from visual arts are influenced by the heterogeneity of the included articles.

**Table S1. The results of the additional meta-analyses for beautiful visual art and beautiful faces**

| Cluster | Volume  (voxels) | Weighted center | | | Maximum  Z-value | Center for maximum Z-value | | | Macroanatomical location |
| --- | --- | --- | --- | --- | --- | --- | --- | --- | --- |
|  |  | x | y | z |  | x | y | z |  |
| ***beautiful > non-beautiful visual art (additional analysis with more studies)*** | | | | | | | | |  |
|  | *No cluster survived* | | |  |  |  |  |  |  |
| ***beautiful > non-beautiful faces (additional analysis with more studies)*** | | | | | | | | |  |
| 1 | 481 | 0 | 48 | -6 | 6.11 | 0 | 48 | -6 | Paracingulate gyrus/Frontal medial cortex |
|  |  |  |  |  | 4.2 | 0 | 36 | 14 | Anterior cingulate gyrus |
|  |  |  |  |  | 3.91 | 0 | 36 | 2 | Anterior cingulate gyrus |
| 2 | 195 | -10 | 16 | -6 | 5.54 | -10 | 16 | -6 | Left accumbens/Caudate |
|  |  |  |  |  | 3.78 | -8 | 10 | -14 | Subcallosal cortex left accumbens |
